# Strain-resolved microbiome sequencing reveals mobile elements that drive bacterial competition on a clinical timescale

**DOI:** 10.1101/125211

**Authors:** Alex Bishara, Eli L. Moss, Ekaterina Tkachenko, Joyce B. Kang, Soumaya Zlitni, Rebecca N. Culver, Tessa M. Andermann, Ziming Weng, Christina Wood, Christine Handy, Hanlee Ji, Serafim Batzoglou, Ami S. Bhatt

**Author notes:** These authors contributed equally to the study (Alex Bishara, Eli L. Moss).

## Abstract

Although shotgun short-read sequencing has facilitated the study of strain-level architecture within complex microbial communities, existing metagenomic approaches often cannot capture structural differences between closely related co-occurring strains. Recent methods, which employ read cloud sequencing and specialized assembly techniques, provide significantly improved genome drafts and show potential to capture these strain-level differences. Here, we apply this read cloud metagenomic approach to longitudinal stool samples from a patient undergoing hematopoietic cell transplantation. The patient’s microbiome is profoundly disrupted and is eventually dominated by *Bacteroides caccae*. Comparative analysis of *B. caccae* genomes obtained using read cloud sequencing together with metagenomic RNA sequencing allows us to predict that particular mobile element integrations result in increased antibiotic resistance, which we further support using *in vitro* antibiotic susceptibility testing. Thus, we find read cloud sequencing to be useful in identifying strain-level differences that underlie differential fitness.

## Introduction

Shotgun short-read sequencing methods facilitate study of the genomic content and strain-level architecture of complex microbial communities. Comparisons of microbial genomes obtained from both isolate sequencing and single-cell techniques have shaped our understanding of strain-level genetic variability^1–3^. Microbial strains of the same species share “core” genes that encode basic phenotypes of the species, but can differ by structural arrangement^4^ or by the presence of accessory genes^5,6^, which facilitate adaptability of the species population as a whole.

Strain-level variation can arise from several mechanisms including horizontal gene transfer and transposon mobilization. Each of these mechanisms is well-described capacity to induce significant changes in phenotype. Through horizontal gene transfer, bacteria can acquire and disseminate genomic elements encoding antibiotic resistance genes, virulence factors, or metabolic capabilities^7,8^. Smaller mobile elements, such as transposons, can affect gene function and regulation by either disrupting coding sequences^9^, or by upregulating neighboring genes through the introduction of strong promoter sequences often carried with the transposon ^10,11^. These transposons can be mobilized during physiological stress, such as exposure to antibiotics, and this mobilization can result in acquisition of improved niche-specific fitness ^12^.

Recent computational techniques using marker gene sets have enabled metagenomic sequencing approaches to be used for the tracking of microbial strains across different samples^13–15^. These methods track strains by first using available isolate references to pre-compute species-specific marker gene sets. Alignments of metagenomic short reads to these markers can then be used to estimate strain-specific single nucleotide variant (SNV) profiles, which can be tracked across samples. Although these approaches provide an efficient solution for the tracking of microbial strains, they are inherently limited to the study of clades with at least one representative isolate reference genome. Furthermore, these methods are unable to distinguish between very closely related strains sharing the same marker SNV profile, but differing in sequence from recent horizontal gene transfer or transposition events.

Dedicated metagenomic assemblers^16–18^, and also binning approaches based on sequence similarity^19–22^ and coverage depth covariance^23,24^, aid analyses of metagenomic short-read sequences without relying on available reference isolates. These techniques can yield significantly more comprehensive draft genomes, but have difficulty in resolving conserved and recently duplicated sequences, such as those arising from recent horizontal gene transfer and transposition. As a result, these regions currently remain unassembled and often unconsidered in analysis using standard short-read metagenomic sequencing. Culture-based methods can be helpful in this context, but have limited throughput and require laborious and often biasing culture steps.

To address the shortcomings of existing metagenomic workflows, we recently developed a metagenomic shotgun sequencing and assembly method that provides complete microbial genomes from complex microbiome samples^25^. Our approach utilizes recent “read cloud” platforms ^25,26^ (also referred to as “linked-reads”^27^), which cost-effectively generate short-read sequences tagged with long-range information. Our novel de novo assembly approach uses this long-range information to assemble the duplicated and conserved sequences that precluded existing workflows from recovering full genome sequences for individual microbial strains.

In this work, we present the first application of read cloud sequencing to longitudinal microbiome samples, and show that it enables the discovery of strain-level sequence variants that confer selective advantages under differing environmental stresses. We apply read cloud sequencing using the 10x Genomics platform and also metagenomic RNA sequencing to study a time series of human stool samples obtained from a patient undergoing hematopoietic cell transplantation treatment for a hematological malignancy. During treatment, the patient received extensive broad-spectrum antimicrobials, imparting profound selective pressure on the intestinal microbiome that resulted in eventual domination by *Bacteroides caccae*. Comparative analysis of the read cloud *B. caccae* genomes and transcriptomes enables us to find overexpression of antibiotic resistance genes coinciding with both antibiotic administration and the appearance of proximal transposons harboring a putative bacterial promoter region. *In vitro* antibiotic susceptibility testing and whole genome sequencing of isolates derived from patient stool samples support functional predictions made using our metagenomic approach.

Our work demonstrates that read clouds can enable culture-free metagenomic sequencing to be used to elucidate sequence variations underlying fitness advantages of microbial strains.

## Results

### Read cloud sequencing of a human intestinal microbiome time series

We applied read cloud sequencing to four longitudinal clinical stool samples obtained from a patient receiving treatment for hematologic malignancy at the Stanford Hospital Blood and Marrow Transplant Unit. Samples were collected over the course of 56 days. The patient was a 46 year old man who underwent hematopoietic cell transplantation (HCT) for myelodysplastic syndrome and myelofibrosis, which was refractory to treatment with azacitidine. The patient received multiple medications during the period of observation, including antibiotics, antivirals, antifungals, chemotherapeutics and immunosuppressives (Figure 1). The patient was diagnosed with gastrointestinal (GI) graft-versus-host disease (GVHD) (clinical grade 3, histological grade 1 in duodenal biopsy) on day 42. The patient received immunosuppressants (tacrolimus, prednisone, prednisolone, budesonide and beclomethasone) between days 7 and 60 as prophylaxis and treatment for GVHD. The patient was on a special low fiber diet, and fermented foods, probiotics, and prebiotics were restricted from the patient’s diet over the entire time of sampling as a part of routine peri-HCT care protocol at this center. On day 43, the patient’s diet was restricted to total parenteral nutrition administered through an intravenous line. This continued for eight days, whereupon the patient began a clear liquid diet. During the course of treatment, the patient’s intestinal microbiome underwent profound simplification, rapidly becoming dominated by *Bacteroides caccae*, a normal resident within a healthy human intestinal microbiome^28^ (Figure 1; genus-level classifications included in Supplementary Figure 1).

**Figure 1.**
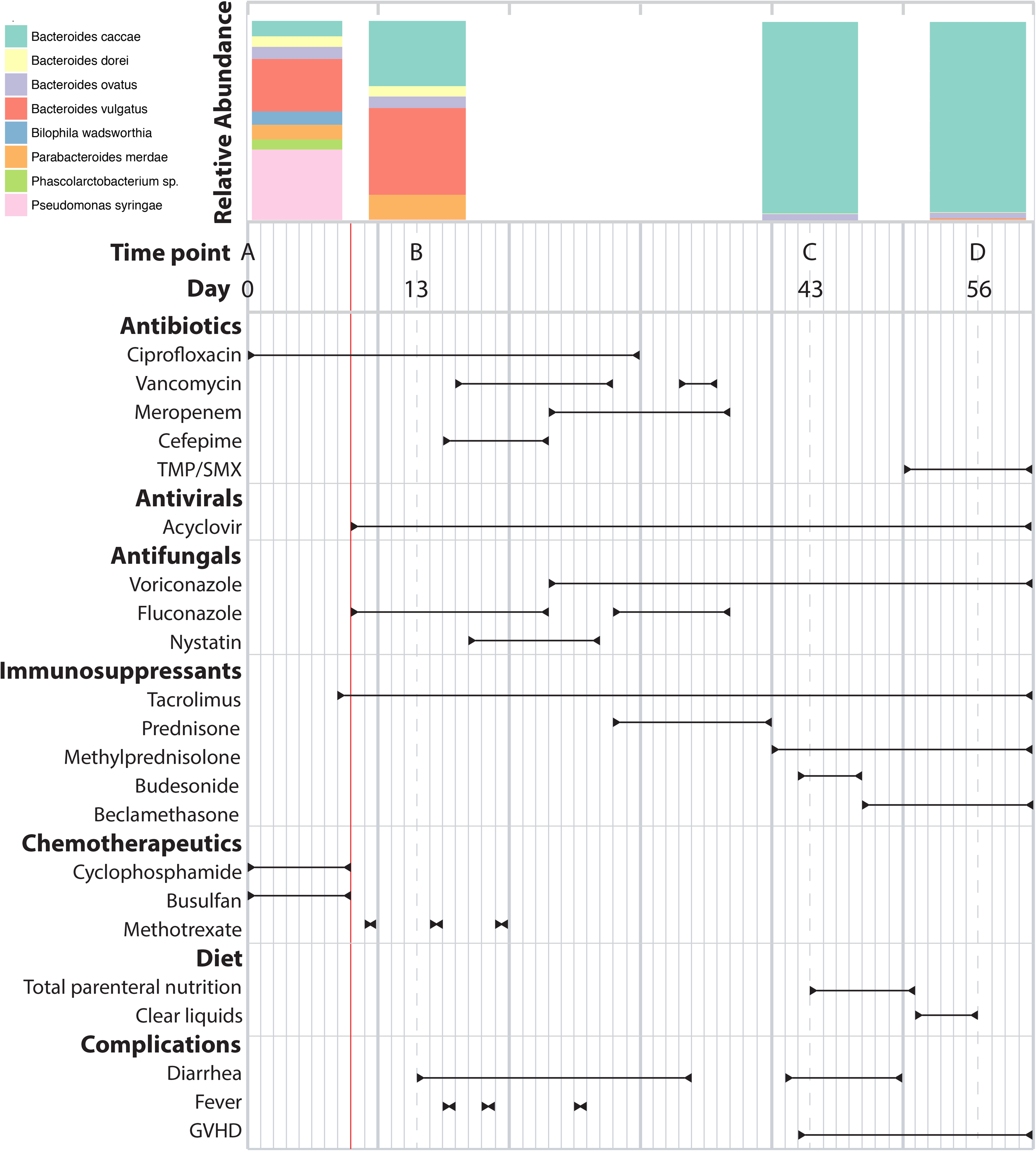
Patient condition, drug exposure, and intestinal microbiome composition during treatment. The study subject was admitted to Stanford Cancer Center with myelodysplastic syndrome and myelofibrosis and subsequently underwent hematopoietic stem cell transplantation (HCT, denoted by a red line). Stool samples were collected prior to HCT and over the following five weeks as the patient underwent chemotherapy, antibiotic treatment, and immunosuppression. Taxonomic classification of sequencing reads (Illumina TruSeq Nano DNA) reveals pronounced dysbiosis emerging following HCT with gut domination by *Bacteroides caccae*, a commensal bacterium. Relative abundances of each species are determined using reads that are classified at the species-level (genus-level classifications shown in Supplementary Figure 2).

To study the trajectory of this patient’s intestinal microbiome throughout treatment, we selected the following four time points for sequencing: **A** (day 0) pre-chemotherapy and pre-HCT; **B** (day 13) post-chemotherapy and post-HCT; **C** (day 43) post broad-spectrum antibiotic exposure and onset of GI GVHD; **D** (day 56) following onset of GVHD and introduction of a new antibiotic regimen including trimethoprim (Figure 1). Read cloud and standard short-read sequencing libraries were prepared from DNA extracted from each of the samples. Additionally, short-read libraries were prepared from RNA extracted from each of the samples (see Methods).

Species-level community compositions obtained by both short-read and read cloud approaches were first assessed using *k*-mer based short-read classifications (Supplementary Figure 1). Both approaches displayed domination by *B. caccae* in later time points C and D, but differed significantly in community composition for earlier time points A and B. Read cloud libraries showed an overrepresentation of Gram-positive bacteria in these time points as compared to short-read libraries. This discrepancy may be the result of differences between the lysis methods used to extract DNA for read cloud and short-read libraries ^29,30^ (see Supplementary Results).

To obtain individual genome drafts for constituent microbes present in each sample, short-read and read cloud libraries were first assembled using either a conventional short-read assembler alone or a short-read assembler and Athena^25^, respectively (see Methods). Contigs in each metagenome draft were then binned and taxonomically classified to obtain annotated genome drafts (see Methods). CheckM^31^ was applied to the resulting bins to assess genome completeness and contamination by the presence of lineage-specific single copy core genes. In addition to assembling *B. caccae* genome drafts, read clouds produced well-assembled genome drafts for other *Bacteroides* members *Bacteroides vulgatus* and *Bacteroides uniformis* (Supplementary Table 2). For time points A and B, read clouds also assembled several high quality genome drafts^32^ with N50 exceeding 500kb for members of enriched Gram-positive genera including *Eubacterium, Lachnospiraceae, Gemella*, and *Flavonifractor*. Read clouds produced single genome bins for *B. caccae* that were both 94% complete and <1% contaminated in dominated time points C and D, but produced multiple, less complete bins for *B. caccae* in earlier time points A and B.

To produce more complete drafts for *B. caccae*, bins annotated as *B. caccae* in each assembly were merged and reevaluated using checkM as more complete genome drafts (Supplementary Table 3). Read clouds with Athena assembly were able to consistently yield more contiguous and more complete drafts of *B. caccae* than short-read sequencing and assembly (Figure 2). Despite differences in overall sequence coverage and community composition, *B. caccae* was determined to have comparable absolute raw sequence coverage in both short-read and read cloud libraries (Supplementary Table 3). Our most contiguous and complete *B. caccae* read cloud draft was from time point C with an N50 of 414kb and total size of 5.5Mb; this is a notable improvement over the best short-read draft, which was from time point C with an N50 of 88kb and total size of 4.7Mb. *B. caccae* coverage varied between 27x and 1542x across read cloud libraries, and 157x and 759x across short-read libraries. All four *B. caccae* read cloud drafts showed large scale structural concordance with the available closed reference genome (Genbank ID GCA_002222615.2), with the exception of one misassembly in the time point D draft around a 16S/23S ribosomal RNA gene operon (Supplementary Figure 2).

**Figure 2.**
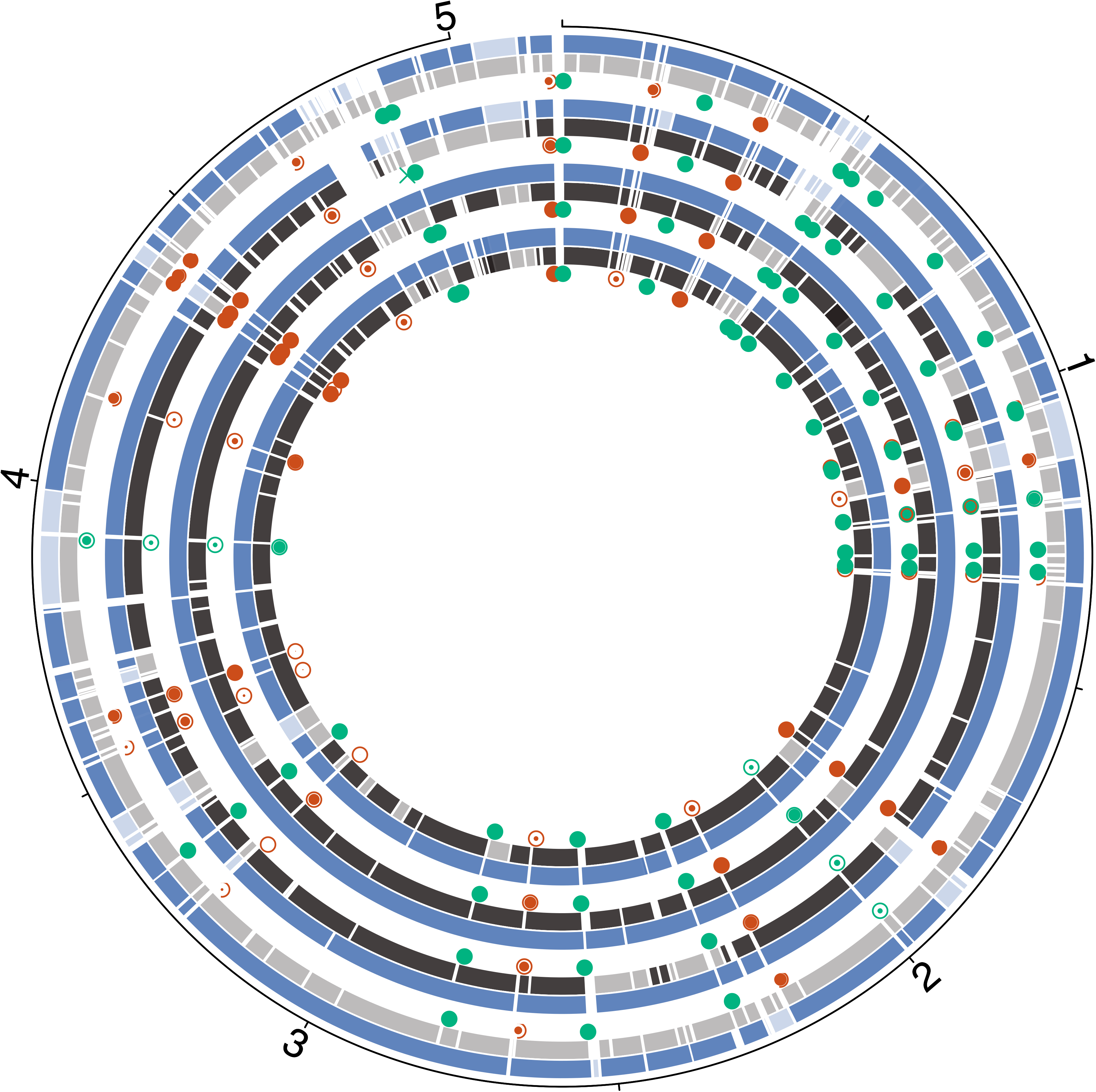
Read cloud and short-read genome drafts of Bacteroides caccae obtained from sequencing of stool samples. Read cloud drafts assembled using Athena (blue) are more contiguous and complete than short-read drafts assembled using conventional assembly (grey). Tracks are ordered chronologically (A through D) from the outermost inward. The read cloud *B. caccae* draft from time point C was the most contiguous and served as the reference for all alignments. Contigs from each read cloud and short-read library are assigned a lighter color if they aligned to this reference, but did not belong to a draft classified as *B. caccae*. Short reads did not produce a draft annotated as *B. caccae* for time point A. The best read cloud and short-read drafts are both obtained through sequencing of time point C (read cloud: 414kb N50, 5.5Mb size, 99.3% complete, 1.6% contaminated; short read: 88kb N50, 4.7Mb size, 97.7% complete, 0.6% contaminated). The read cloud drafts include a total of 18 assembled integration sites of IS614 (red circles) and 25 assembled integration sites of a candidate insertion sequence (green circles) that are missing from all short-read drafts. Alignments of raw short reads to these sites indicated the presence of both strains without insertion sequence integration and strains with the insertion sequence integration. Estimated proportions of these strains for each site and time point are shown with different filled in areas of each circle, with an empty circle denoting predominance of ancestral strains lacking an IS at that location and shaded circles denoting predominance of strains with the IS at that location.

We next compared the best read cloud and short-read *B. caccae* drafts, which were both produced in libraries from time point C, to available reference isolate genomes. We first aligned the six available reference isolate genomes of *B. caccae* (Genbank IDs NZ_AAVM02000021.1, NZ_JH724079.1, NZ_CZBL01000001.1, NZ_CZAI01000001.1, NZ_CP022412.2, NZ_PUEQ01000001.1) to our read cloud drafts using MUMMER^33^. The read cloud *B. caccae* genome contained 639kb of novel sequence not represented in any reference isolate, compared to just 318kb of novel sequence contained in the short-read draft. The median sequence identity for alignable bases between our *B. caccae* drafts and the reference isolates was 99.5% for both read clouds and short reads (Supplementary Table 4).

### Read clouds recover nearly identical strains in clinical samples

We posited that comparative analysis of the *B. caccae* genome drafts across time points may yield insight into either selection or potential genomic remodeling that this organism may have undergone as it grew to eventually dominate this host’s intestinal microbiome.

We first searched the read cloud *B. caccae* drafts for differing large-scale genomic island incorporations. Pairwise alignments of *B. caccae* drafts from successive time points were first obtained with MUMMER^33^, and we used these alignments to identify large genomic sequences that were assembled into differing genomic contexts between drafts of different time points (see Methods, Supplementary Figure 3). We identified a total of five separate genomic islands ranging from 17kb to 71kb in size that were integrated into different genomic contexts between drafts of different time points (Supplementary Table 5). Two of these islands were also present in the draft genomes of *B. vulgatus* and *B. uniformis*.

We next searched the read cloud *B. caccae* drafts for small-scale insertion sequences. Insertion sequence elements were first identified by counting *k*-mers in each read cloud *B. caccae* draft, selecting high frequency *k*-mers, and assembling these to determine precise sequence (see Methods). This procedure yielded two putative insertion sequence elements. The first IS, 1596bp in length, was annotated using nucleotide alignments with BLAST (nt database)^34^ It was determined to be IS614, a conserved Bacteroides insertion sequence (IS). IS614 encodes a transposase as well as an outward-facing promoter sequence, which has been predicted to drive transcription of genes neighboring the IS^10^. The second IS, 1470bp in length, could not be annotated as a previously described IS, but shares protein sequence homology and a conserved DDE domain with the IS4 insertion sequence family. Both IS614 and the unannotated candidate IS appear in the metagenome drafts of the short-read libraries, but appear only in single copies detached from genomic context with extreme sequence coverage depth (16,664x and 14,615x coverage for IS614 and the candidate IS in time point C, respectively). This suggests that high copy sequence elements, including these insertion sequences, greatly limit the overall quality of genome drafts obtained using standard short-read assembly. Although we found the candidate IS to be present within two reference isolate genomes for *B. caccae* (Genbank IDs: NZ_CZBL01000001.1, NZ_CZAI01000001.1), we could not identify IS614 in any of the six available reference isolate genomes.

We searched for integrations of IS614 and the candidate IS across all read cloud metagenome drafts in which at least 3kb of flanking sequence were assembled on both sides of the integration. We found 28 unique integrations of IS614, 18 of which were determined to be in *B. caccae*, and the other 10 of which were within contigs classified as *B. vulgatus*. We found 25 unique integrations of the unassigned IS, all of which were determined to be in *B. caccae*.

At many insertion sites in *B. caccae*, alignments of the short-read data to the Athena assembly confirmed the co-occurrence of both strains harboring the IS and strains with the pre-insertion ancestral sequence (Figure 3a). From these alignments, we obtained an estimate of the relative abundance of ancestral and insertion-containing strains for each site (see Methods). Only three of the 25 identified integration sites of the candidate IS had short-read alignments indicative of a pre-insertion ancestral sequence. The rest of these integrations appeared to be fixed in the *B. caccae* strain population. In contrast, we observed large shifts in ancestral abundance at several *B. caccae* IS614 integration loci (Figure 3b, Supplementary Table 6), with 15 large (>30%) shifts occurring between consecutive time points.

**Figure 3.**
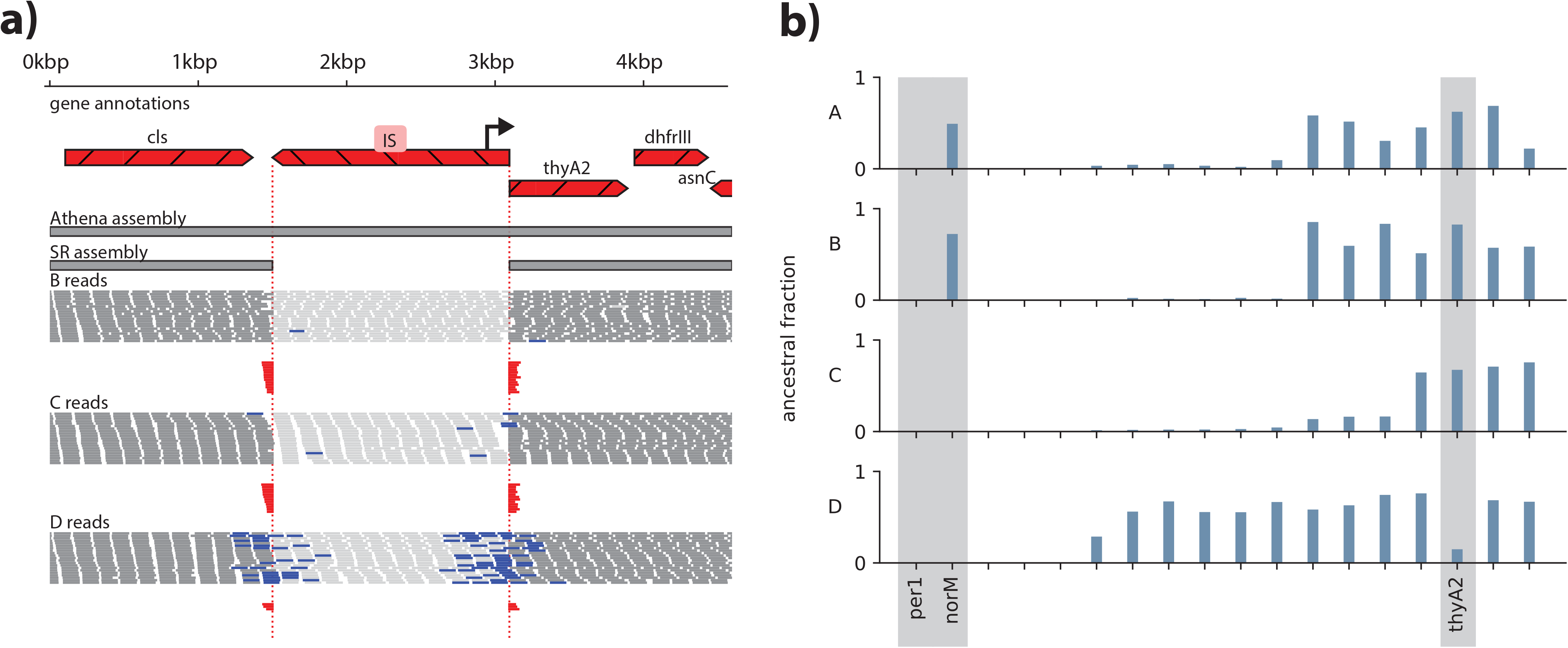
Co-occurrence of multiple B. caccae strains with differing IS614 integrations. a. Alignments of short reads from time points B, C, and D to a representative IS integration site reveal domination of the strain without the IS (“ancestral strain”) in time points in B and C, and domination of the strain harboring the insertion in D. Short-read alignments from B and C show many reads spanning over both left and right junctions (red, indicating global alignment to the ancestral sequence), while short-read alignments from D show many reads supporting the IS integration (blue, indicating read pairs or single reads spanning into the IS). This demonstrates that the IS is present at this locus at time point D but undetectable at timepoints B and C.
b. Estimated relative abundances of *B. caccae* ancestral strains and strains with an IS integration for all 18 detected IS614 integration sites. Integrations upstream of annotated antibiotic resistance genes *norM, thyA2*, and *per1* are shaded. Major shifts in abundances amongst the strains with and without integrations upstream of *norM* and *thyA2* can be seen between time points B and C and time points C and D, respectively. Integration sites are sorted by ancestral strain fraction in time point C.

### Identification of insertion-mediated transcriptional upregulation

We next applied RNA sequencing at each time point to investigate the potential transcriptional effects of the genomic alterations we detected in *B. caccae*, focusing on IS614. This IS contains a putative outward-facing promoter near its 5’ end oriented antisense to its transposase coding sequence^10^. Determining the transcriptional effect of this IS is difficult in a complex metagenomic setting, as RNA sequencing reads may originate from co-occurring strains with or without a given insertion. In light of this difficulty, we restricted our attention to integration sites that were dominated first by ancestral strains and then by IS-harboring strains in consecutive time points, with at least 30% change in estimated ancestral abundance. In these sites, a corresponding increase in transcription of genes downstream of the promoter versus upstream of the promoter is more likely attributable to the additional promoter introduced by the IS. We found five such candidate loci, four of which had putative promoter sites introduced in the same sense as the adjacent downstream gene. All four of these loci showed “transcriptional asymmetry” in which transcription of the downstream neighboring gene was increased between three-fold and eight-fold relative to transcription of the upstream neighboring gene coinciding with an increase in relative abundance of strains harboring the insertion (Figure 4, Supplementary Table 7).

**Figure 4.**
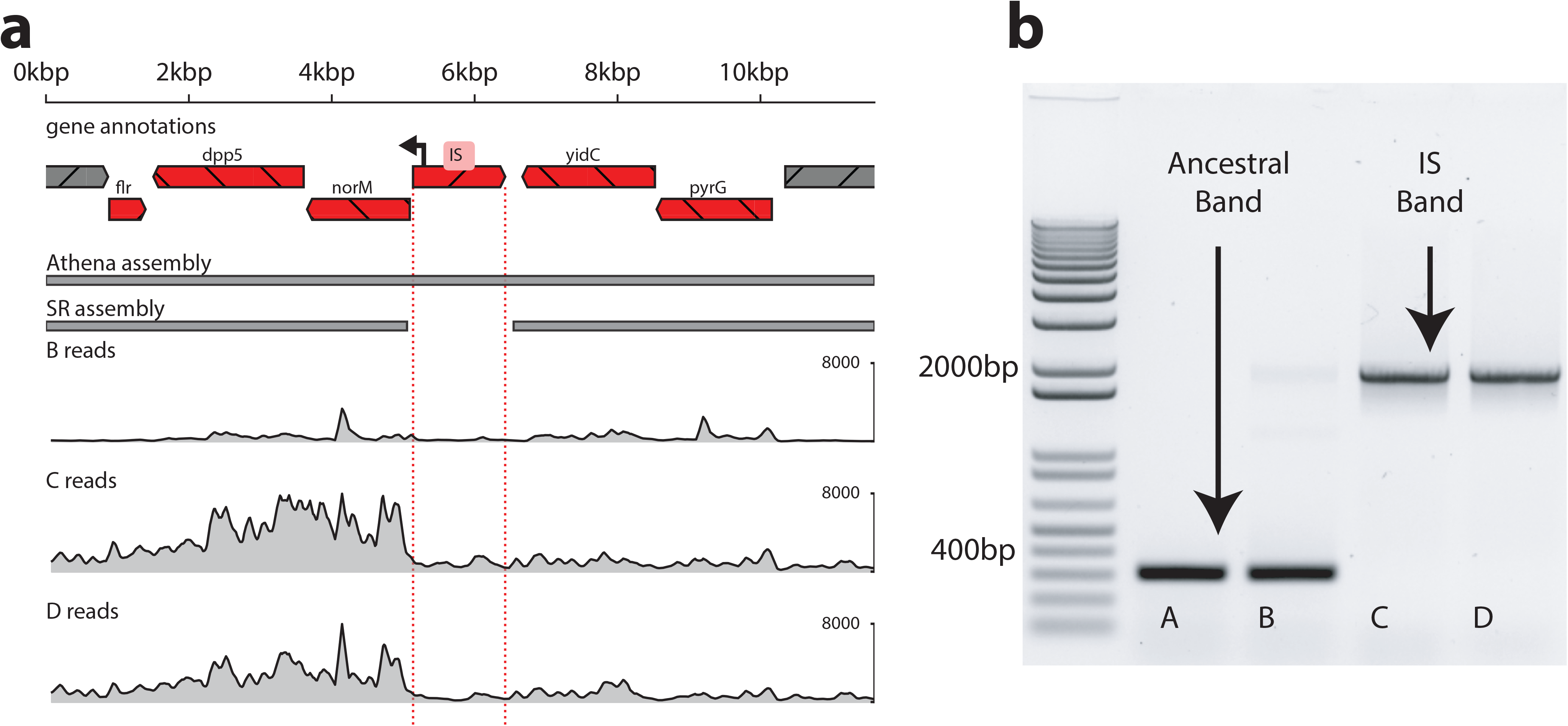
Metagenomic RNA sequencing supports IS-mediated transcription within B. caccae. IS614 contains a putative promoter that may affect transcription of neighboring genes. Comparison of RNA sequencing read depths of genes upstream and downstream from this promoter can give insight into its relative contribution to transcription. In later time points C and D, dominant strains harbor an introduced promoter positioned to upregulate *norM*. In prior time point B, dominant strains have no IS in this region.

a. In time point B, which is dominated by ancestral strains without the promoter, RNA sequencing read coverage depth is relatively equal on both sides of the integration site. In time point C, which is dominated by strains with the introduced promoter, the coverage of the downstream gene *norM* is much higher than the upstream gene *yidC*. The coverage depths of all neighboring genes increases in time point C relative to B, but this increase is 10-fold greater in genes downstream of the introduced promoter. Read pairs spanning between the IS promoter and *norM* further support the transcriptional contribution of the IS. This difference in coverage and domination by strains with this promoter both persist through time point D.
b. PCR with primers flanking the above integration instance of IS614 yields amplicons without the insertion sequence in earlier time points A and B (400bp), and with the insertion in later time points C and D (1.9kb).

One notable transcriptional asymmetry coincided with placement of the putative promoter in IS614 to upregulate *norM*, a multidrug resistance transporter (Figure 4a). *NorM* is a multidrug efflux protein that can confer resistance to ciprofloxacin^35^. Ciprofloxacin was administered for the first 30 days of treatment through time points A and B (Figure 1). Manual inspection of short-read alignments to this insertion site showed this integration to be undetectable in time point A, present in roughly a third of strains in B, and then in the majority in time points C and D, consistent with visible band patterns in our targeted PCR results (Figure 4b, Supplementary Table 6). Another transcriptional asymmetry was observed adjacent to *thyA2* and *dhfrIII*, encoding thymidylate synthetase and dihydrofolate reductase, respectively (Supplementary Figure 4). Both are linked to trimethoprim sensitivity^36,37^, and the marked rise in strains carrying an adjacent IS-borne promoter coincides with the administration of this antibiotic to the patient prior to the final time point D. A third transcriptional asymmetry was found in resA, an oxidoreductase involved in cytochrome c synthesis^38^ (Supplementary Figures 5). The abrupt changes observed in the abundance of insertion sequences adjacent to these loci suggest selective pressures applied to the bacterial strains.

We found the most highly expressed gene in time points C and D to be the extended-spectrum beta-lactamase gene *per1*, known to confer resistance to beta lactam antibiotics^39^, which were administered to the patient between time points B and C. *Per1* was expressed nearly 60% more than the second most expressed gene in both time points C and D. (see Supplementary Results)

Though several insertion sequences became undetectable in DNA sequence data between timepoints C and D, strains with insertions adjacent to *norM* and *per1* continue to dominate through the end of the investigated time course. By time point D, ciprofloxacin and meropenem have been withdrawn for 26 and 19 days, respectively, yet expression of resistance genes *norM* and *per1* remained increased compared to levels prior to antibiotic exposure.

### *B. caccae* isolate sequencing and antibiotic susceptibility testing

To assess phenotypic predictions made by evaluating the *B. caccae* genome drafts generated by our read cloud approach, we isolated *B. caccae* strains from stool samples for whole genome sequencing and antibiotic susceptibility testing. Stool samples from each of the four time points A, B, C, and D were streaked directly onto culture plates with selective media in order to isolate members from the *Bacteroides fragilis* group including *B. caccae* (see Methods). A total of 53 colonies from time points A, C, and D were selected based on morphology and subsequently cultured in liquid media. We were unable to obtain colonies representative of the *B. fragilis* group using culture plates from the time point B stool sample. The resulting isolates were sequenced and assembled to obtain draft genomes (see Methods, Supplementary Table 8).

Analysis of the assembled genomes revealed 14 of the 53 isolates were determined to be an organism other than *B. caccae* (Supplementary Table 8). Notably, 7 of 10 of our time point A isolates were determined to belong to another *Bacteroides* species closest to *Bacteroides uniformis*. Our isolate collection was determined to contain a total of 41 *B. caccae* strains with three, 17, and 21 *B. caccae* strains from time points A, C, and D respectively.

We examined sequencing data from the 41 *B. caccae* isolates to discover genomic alterations that may potentially be under selection between different time points over the course of the patient’s treatment. We first detected the set of all flanking genomic sequences that were downstream of the putative bacterial promoter present in IS614, and also larger genomic islands that were differentially present between the isolates (see Methods). Each *B. caccae* isolate was then genotyped for the presence or absence of each IS614 integration and genomic island incorporation.

Four IS614 integrations were determined to be present in all *B. caccae* isolates, including *per1*, which we predicted to be fixed in the *B. caccae* strain population across time points using our read cloud metagenomic approach. Hierarchical clustering of *B. caccae* isolates by IS614 integrations revealed the presence of four distinct strain subpopulations that shift in abundance between time points (Figure 5). These observed shifts in strain subpopulation abundance are concordant with predictions made using our read cloud metagenomic approach. One observation that was enabled by isolate sequencing and was missed by our original read cloud genome assemblies was the identification of an IS614 repeat present upstream of the gene *susC*, a surface-accessible protein involved in starch binding and utilization ^40^. This IS614 instance was present in high abundance in the latter time point isolates (15 out of 17 time point C isolates, 8 out of 21 time point D isolates), concordant with the time period of poor oral dietary intake and initiation of total parenteral nutrition.

**Figure 5.**
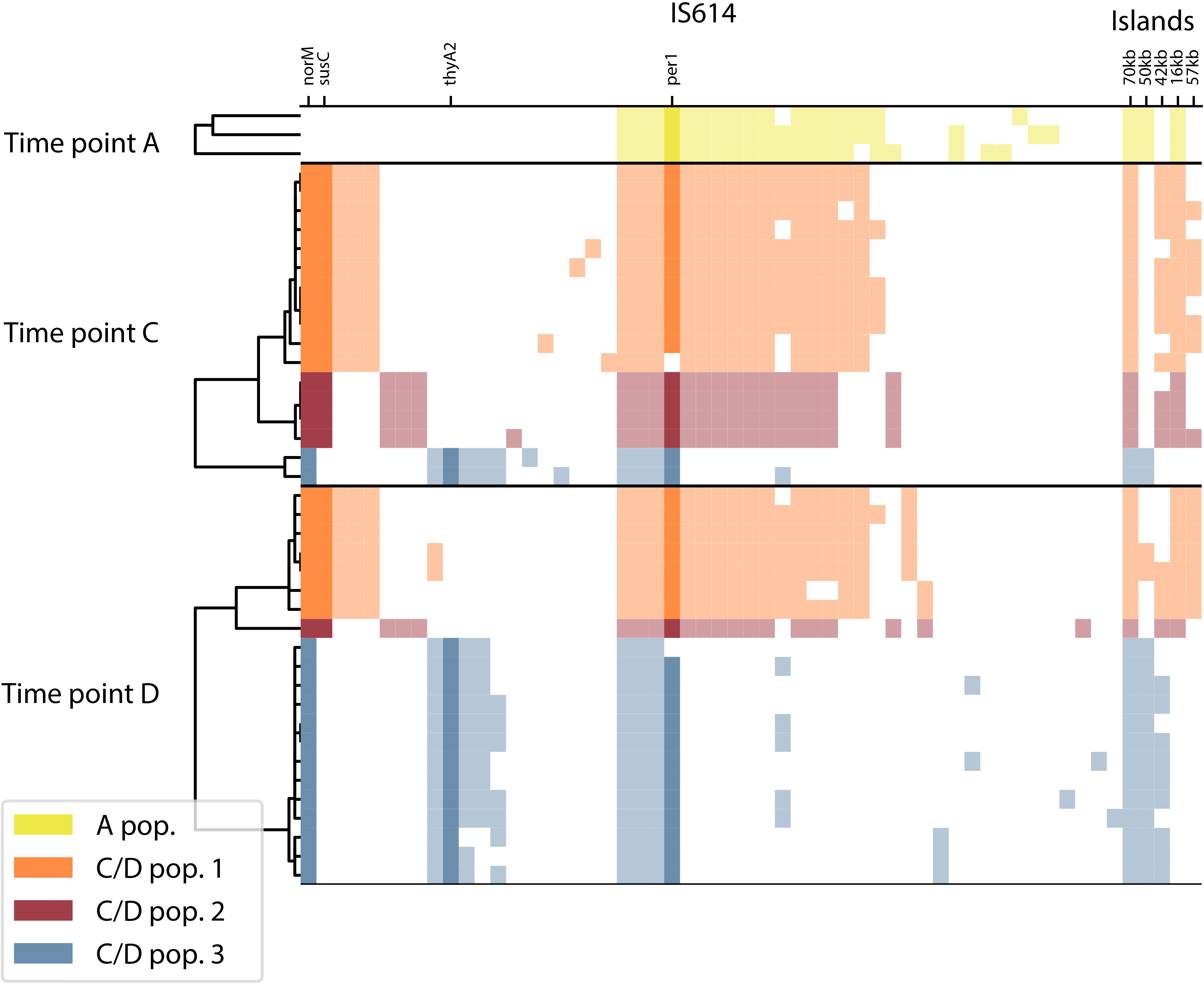
Genomic alterations detected in B. caccae isolates supports selection throughout treatment. The detected proportions of *B. caccae* isolates (rows) with different IS614 integrations and large-scale genomic island integrations (columns; filled squares indicates presence) are concordant with estimated proportions obtained using read clouds. Hierarchical clustering of isolates by their IS614 integrations within each time point reveals three distinct subpopulations of *B. caccae* strains, which appear to shift in relative abundance between time points C and D. The IS614 integration upstream of *norM* is the only one that is absent from all isolates from time point A (before ciprofloxacin exposure), but appears in all isolates from time point C and D (after ciprofloxacin exposure). The IS614 integration upstream of *per1* was determined to be present in all of the isolate *B. caccae* strains we obtained. Initial analysis of the isolate sequencing data was unable to detect IS614 integrations in front of *per1* for two isolates, but manual inspection of the assembly graphs of these confirmed this integration to be present in these two as well (outlined squares). The IS614 integration upstream of *thyA2* appears in the minority of strains in time point C (before trimethoprim exposure), and appears in the majority of strains in time point D (after trimethoprim exposure). The IS614 integration upstream of *susC*, which was detected in isolate sequence data, also appears to be under selection in time point C. Unlike the IS614 integrations, none of the large-scale genomic islands appear to be under selection between time points.

All strains from time points C and D (post ciprofloxacin exposure) contain an IS614 integration upstream of *norM*, which is absent from all three strains from time point A (pre ciprofloxacin exposure). Although a minority of strains with an IS614 integration upstream of *thyA2* and *dhfrIII* are present in time point C (pre trimethoprim exposure), the majority of strains with this integration are present in time point D (post trimethoprim exposure). The observed shifts in abundance of the *B. caccae* strain subpopulations are consistent with the differing selective pressures applied from the patient’s drug regimen over the course of treatment. Notably, while clear subpopulations with varying IS614 positions appear to exist and have a selective fitness advantage at various time points, the larger genomic islands appear to be sporadically integrated amongst the individual strains. This suggests that none of the larger genomic islands appeared to be under selection between time points A and C or between time points C and D.

We next tested the antibiotic susceptibility of several representative strains from each *B. caccae* subpopulation to determine if the identified strain-level variants resulted in differing resistance phenotypes. All three strains from time point A and four strains from each of the three subpopulations in time points C and D were selected for susceptibility testing with ciprofloxacin and trimethoprim (Figure 6a). Each selected isolate strain was tested against different concentrations of each drug using between two to four replicates with a broth microdilution Minimum Inhibitory Concentration (MIC) method adapted from Clinical Laboratory Standards Institute (CLSI; see Methods). Growth curves were generated for each isolate, in replicate, by measuring optical density at a wavelength of 600 nm (OD600) over a range of drug concentrations (Supplementary Figures 6, 7). Area-under-the-curve (AUC) analysis of these OD600 growth curves allowed determination of MICs for each replicate (see Methods).

**Figure 6.**
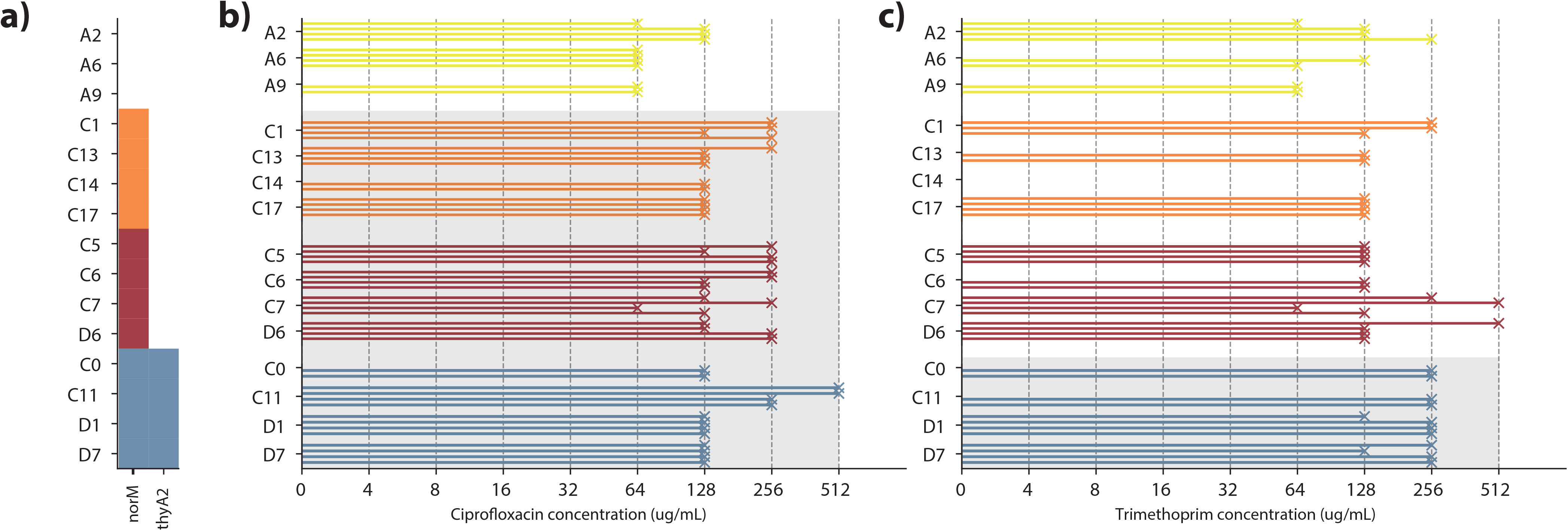
Antibiotic susceptibility testing of B. caccae isolates from each strain population. All three *B. caccae* isolates from time point A and four *B. caccae* isolates from each subpopulation present in time points C and D (15 isolates in total), were tested for their susceptibility to antibiotics ciprofloxacin and trimethoprim as received by the patient during treatment. Each isolate was tested against each antibiotic using two to four replicates. Replicates showing minimal growth in the absence of antibiotic exposure were removed from consideration in determining minimum inhibitory concentrations (MICs).

a. The presence (filled square) of different IS614 integrations within tested isolate strains that are upstream of annotated genes *norM* and *thyA2*, which may affect resistance to ciprofloxacin and trimethoprim. These integrations result in introduction of likely bacterial promoters upstream of these genes likely leading to increased expression and as a result, increased antibiotic resistance. *NorM* is a known multidrug efflux pump that can confer resistance to ciprofloxacin. Upregulation of *thyA2/dhfrIII* has been shown to affect resistance to trimethoprim^37^.
b. MICs of *B. caccae* strains for ciprofloxacin. Strains from time points C and D with IS614 integrations predicted to increase resistance to ciprofloxacin (shaded) show markedly higher MICs (2-4 fold) than strains from time point A.
c. MICs of *B. caccae* strains for trimethoprim. Strains from all subpopulations showed high resistance to trimethoprim. Strains from the third subpopulation in time points C and D with IS614 integrations predicted to increase resistance to trimethoprim (shaded) show a modest increase in resistance as compared to other subpopulations.

Susceptibility testing of selected isolates revealed the overall *B. caccae* strain population to be highly resistant to both ciprofloxacin and trimethoprim, with varying degrees of resistance observed between isolated strains from the four subpopulations (Figure 6). Previously reported MIC50 values (lowest concentration of the antibiotic at which 50% of the tested isolates were inhibited) for members of the *B. fragilis* group have been ≤ 8 ug/mL for ciprofloxacin^41,42^ and ≤ 8.8 ug/mL for trimethoprim ^43^. By contrast, our observed MIC50 values across all subpopulations were ≥ 64 ug/mL for both ciprofloxacin and trimethoprim. Nonetheless, varying degrees of the drug resistance phenotypes were observed between isolated strains from the four subpopulations. Isolate strains from time points C and D, which were predicted to be more resistant to ciprofloxacin using our metagenomic approach, showed higher MICs (two-fold to four-fold) than isolate strains from time point A (Figure 6b). Isolate strains from the third subpopulation present in time points C and D, which we predicted to be more resistant to trimethoprim, showed a more modest increase (less than two-fold) than isolate strains from the other subpopulations (Figure 6c). Grouping of replicates by subpopulation assignment enabled the examination their collective growth under varying drug concentrations, and the observed drug sensitivities were largely consistent with the determined MICs (see Supplementary Figure 8 and Methods).

Whole genome sequencing of *B. caccae* isolate strains from time points A, C, and D together with antibiotic susceptibility testing support the presence of selection amongst strain populations of *B. caccae* with different IS614 integrations as predicted using our read cloud metagenomic approach.

## Discussion

We present an application of a recent read cloud metagenomic sequencing approach to a clinical microbiome time series. Using this method, we are able to resolve differential positioning of mobile elements within microbial strains present in the patient’s intestinal microbiome over the course of treatment. Read cloud genome drafts together with RNA sequencing allow the detection of mobile element-mediated transcription of antibiotic resistance loci, suggesting potential differential antibiotic resistance between strains. Whole genome sequencing and *in vitro* antibiotic susceptibility testing of microbial strains isolated directly from patient stool samples provide compelling evidence to support these predictions. Our work shows that culture-free metagenomic sequencing approaches can be used to investigate adaptive regulatory variations mediated by mobile elements, and demonstrates their involvement in clinical timescale strain dynamics within the gut microbiome.

The clinical subject we examine in this study is an individual who underwent extensive treatment with several classes of medication while undergoing hematopoietic cell transplantation. Immune suppression and extensive antibiotic treatment likely rendered this individual highly susceptible to destabilization of intestinal microbiome. In other studies, this type of poorly diverse intestinal microbiota has been associated with increased overall mortality, gastrointestinal graft-versus-host disease GI GVHD and other complications^44–47^. Previous studies of similar cohorts as well as healthy individuals have shown that antibiotic use can lead to intestinal domination by one or few microorganisms^48,49^, but the mechanisms by which any specific microorganism achieves dominance over the larger community remain incompletely understood. Our study of this patient revealed that populations of microbes with apparently stable taxonomic composition can, in actuality, be composed of many closely related strains that undergo wide fluctuations in abundance.

Using our read cloud sequencing and assembly approach, we observe large shifts abundance between bacterial strains harboring different IS614 integration sites. Through examination of RNA sequencing data, we find evidence of insertion sequence mediated transcription of genes downstream of these integration sites. One IS614 integration site positioned to upregulate *norM* rises was found to be undetectable in *B. caccae* strains within time point A, present in a fraction of strains within time point B, and present in all detectable strains within time points C and D. This rise to dominance is consistent with both the observed timing of ciprofloxacin administration and described antibiotic specificity of *norM*^35^. Another IS614 integration site positioned to upregulate *thyA2* and *dfhrIII* is detected in strains at low abundance within time point C, and present in the majority of detectable strains in time point D. This observed shift in strain abundance is again consistent with trimethoprim administration between time points C and D, as well as previous reports associating the increased expression level of these genes with trimethoprim resistance^36,37^. Examination of isolate sequencing data revealed an additional integration positioned to upregulate *susC*, a gene involved in starch binding and utilization^40^. In light of the restrictions placed on the patient’s diet starting after time point B, it is possible that upregulation of *susC* may have rendered organisms with this selective advantage more successful in competing for limited starches available within the gut lumen. Our results suggest that insertion sequences can mediate changes in gene transcription within individual strains, and that the pool of phenotypic variation this creates may allow adaptation to changing environmental stresses.

Our results highlight the importance of *de novo* characterization of microbial communities in order to capture strain-level variation, as well as the importance of strain dynamics in antibiotic-associated dysbiosis of the gut microbiome. Previous methods to detect genomic strain diversity rely on compiled reference sequence collections, and characterize nucleotide divergence within either predefined gene sets or gene presence within a predefined organism-specific pan-genome^50–52^. Although these methods differ substantially, their shared reliance on the reference sequence collection restricts sensitivity in the context of poorly sequenced or unknown species, comprising a large fraction of microbial diversity^53^, or previously unobserved structural variants of known species. Existing short-read methodologies fail to assemble certain classes of sequence, including insertion sequences and larger scale genomic island incorporations, both of which can impart significant changes to bacterial phenotypes. Our results demonstrate the importance of generating high quality individual genome drafts in order to fully understand the complex interactions between strains and species within the microbiome.

Further investigation is needed to determine if the rise in abundance of strains carrying different insertion sequence integrations is due to selection or an active biological process occurring in response to environmental stress. Although strains carrying the IS614 integration upstream of *norM* are undetectable by our approach or PCR in time point A before ciprofloxacin administration, it is possible that these strains were present at very low abundance and grew to dominate in later time points due to selection. Further *in vitro* adaptive evolution experiments that mimic the harsh conditions present in this patient’s can be used to determine whether IS614 is capable of mobilizing in response to these conditions to actively generate strain diversity. Whole genome sequencing of *B. caccae* strains isolated directly from the patient’s stool samples indicates the presence of four major strain subpopulations. Three of the four subpopulations all contain this particular *IS614* integration, which suggests potential convergent evolution. However, a higher number of isolate genomes from earlier time points and also examination of more individual patients would provide more compelling evidence to support this claim.

As methods for read cloud and long read sequencing mature, we anticipate that applications of these methods to longitudinal microbiome samples will illuminate mobile element diversity within other complex microbial communities, and enable further investigation into particular elements that may influence fitness under different environmental stresses.

## Methods

### Clinical participant recruitment

A hematopoietic cell transplantation patient was recruited at the Stanford Hospital Blood and Marrow Transplant Unit under an IRB-approved protocol (Principal Investigator: Dr. David Miklos, co-Investigator: Dr. Ami Bhatt). Informed consent was obtained. A comprehensive chart review was carried out to identify clinical features of the patient, duration and exposure to medications, and diet.

### Sample Collection

Stool samples were obtained from the study subject on an approximately biweekly basis, when available. Stool samples were placed at 4°C immediately upon collection, and processed for storage at −80°C the same day. Stool samples were aliquoted into 2mL cryovial tubes with either no preservative or 700μL of RNAlater and homogenized by brief vortexing. The aliquots were stored at −80°C until extraction.

### DNA library preparation

Stool DNA was extracted for short-read libraries and Gemcode read cloud libraries with the QiAMP Stool Mini Kit (Qiagen, Hilden, Germany) modified with an additional step after addition of ASL buffer consisting of 7 cycles of alternating 30 second periods of beating with 1.0mm Zirconia/Silica beads in a Mini-Beadbeater (Biospec Products, Bartlesville, OK) and chilling on ice. Stool DNA was extracted for Chromium read cloud libraries with the Gentra Puregene Yeast/Bacteria kit (Qiagen, Hilden, Germany), modified with a chilling step at −80°C for five minutes, followed by ethanol DNA precipitation at 14,000g for 20 minutes at 4°C.

Prior to read cloud sequencing library preparation, DNA was size selected with the BluePippin instrument (Sage Science, Beverly, MA). A 5kb-50kb size range was used for Gemcode libraries, and a 10kb-50kb size range for Chromium libraries. Read cloud libraries were then prepared with either the Gemcode or Chromium instrument (10X Genomics, Pleasanton, CA).

DNA used for short-read library preparation was not size selected. Short-read libraries were prepared with the Truseq DNA HT library prep kit (Illumina, San Diego, CA).

All library fragment sizes were assessed with the 2100 Bioanalyzer instrument (Agilent Technologies, Santa Clara, CA) using the High Sensitivity DNA chip and reagent kit. DNA and library concentration estimations were performed using fluorometric quantitation with the Qubit 3.0 fluorometer using the Quibit dsDNA HS kit (ThermoFisher Scientific, Waltham, MA).

### RNA library preparation

RNA was extracted with the Qiagen RNeasy Mini kit from stool samples stored in RNAlater at −80°C. Original total RNA concentration was assayed with the Qubit RNA HS kit. RNA was then ethanol precipitated and resuspended in nuclease-free water to concentrate, and then quantified again using both Qubit RNA HS and Qubit DNA HS kits to determine the degree of DNA contamination. Contaminating DNA was removed using the Baseline-ZERO DNase protocol (Epicentre, Madison, WI) with 30 minute incubation followed by a second ethanol precipitation. Ribosomal RNA was depleted with the Epicentre Ribo-Zero rRNA removal kit (Bacteria) and purified with another ethanol precipitation. The rRNA-depleted mRNA quality was assessed with the 2100 Bioanalyzer using the Agilent RNA6000 Pico kit and quantified with Quibit RNA HS assay. cDNA sequencing libraries were then prepared with the Illumina Truseq Stranded mRNA kit following the Truseq Stranded mRNA LT protocol. Resulting DNA libraries were quantified with Qubit DNA HS kit and their quality evaluated by 2100 Bioanalyzer instrument using the High Sensitivity DNA chip and reagent kit.

### Sequencing

Truseq libraries were sequenced with 2×101bp reads on an Illumina HiSeq 4000 instrument, each library receiving 2-6Gb sequence coverage with the exception of time point A, which was additionally sequenced with 34Gb of coverage after the initial attempt produced insufficient coverage. 10x Gemcode libraries were sequenced with 2×148bp reads on an Illumina NextSeq 500 instrument, with each library receiving 4-7Gb sequence coverage. 10x Chromium libraries were sequenced with 2×151bp reads on one lane of Illumina HiSeq 4000, each receiving 18-22Gb of sequence coverage (Supplementary Table 2). RNAseq libraries were sequenced with 2×101 bp reads on an Illumina HiSeq 4000 instrument, each library receiving 8-12Gb sequence coverage with the exception of time point A, for which a high quality RNA sequencing library could not be obtained.

### Metagenome assembly and genome draft generation

Raw reads from all DNA libraries were first subjected to the same quality control and trimming as follows. Sequence data were trimmed using cutadapt^54^ using a minimum length of 60bp and minimum terminal base score of 20. Reads were synced and orphans (reads whose pair mates were filtered out) were placed in a separate single-ended fastq file with an in-house script. RNA sequencing reads displayed uniform high quality and were not trimmed.

Trimmed reads from all libraries were then assembled using metaSPAdes 3.11.1 ^55^ with default parameters for paired-end input. MetaSPAdes seed assemblies obtained from read cloud libraries were then further assembled using Athena^25^. Assemblies were visualized with IGV^56^, R^57^ and python using the ggplot2^58^, circlize^59^ and matplotlib^60^ libraries.

For read cloud and short-read libraries, coverage was calculated for assembled contigs by aligning raw short reads with BWA v0.7.10^61^. Metabat v2.12.1 ^62^ was then used to group contigs to form draft genomes. Drafts were then evaluated for a number of criteria to assess quality: Metaquast v4.6.0^63^ for N50 and assembly size, CheckM v1.0.7 ^64^ for genomic completeness and contamination, Prokka v1.12 ^65^ for gene counts, Aragorn v1.2.36^66^ for tRNA counts, and Barrnap v0.7 ^67^ for rRNA subunit counts. Drafts were denoted “high quality” when they contained 18 or more tRNA loci, at least one occurrence each of the 5S, 16S and 23S ribosomal RNA subunits, and achieved a checkM score of at least 90% completeness and at most 5% contamination in accordance with existing standards^68^. Drafts were otherwise denoted “complete” if they achieve the same completeness and contamination criteria. All other drafts were denoted “incomplete.”

Individual contigs from all assemblies were assigned taxonomic classifications using Kraken v0.10.6^69^ with a custom database constructed from the Refseq and Genbank^70,71^ bacterial genome collections. Genome drafts were given taxonomic assignments with a consensus approach: a draft received a species assignment if 60% or more of total bases shared the species-level classification. Drafts were otherwise assigned the majority genus-level classification.

### Discovery of genomic island integration and insertion sequence loci in read cloud metagenomic drafts

To discover large-scale genomic island incorporations, we first obtained pairwise sequence alignments of *B. caccae* genome drafts between time points (A, B), (B, C) and (C, D) using MUMMER^33^. We searched these alignments for instances in which a single contig from one time point produced a gapped alignment over a single contig from another time point, spanning a putative genomic island (Supplementary Figure 3). Potential genomic island incorporations that were not assembled within a single contig could not be fully resolved and were not considered. No differential genomic island incorporations were observed in time point B with respect to time point A. Three separate genomic islands (17kb, 57kb, and 71kb) were found in the *B. caccae* genome within time point C, but were absent from the draft in time point B. The 17kb and 57kb islands were also found within the draft genomes of *B. vulgatus* and *B. uniformis*. Two genomic island incorporations of sizes 43kb and 50kb were observed in time point D with respect to time point C. Neither of these two islands were assembled into alternate genomic contexts in time point C.

To discover smaller-scale insertion sequences, we examined high frequency *k*-mer sequences that were present in the *B. caccae* genome. We first obtained *k*-mer counts from the time point C read cloud draft using Jellyfish^72^ with k = 101. The vast majority of *k*-mers originate from single copy portions of the genome, but we isolated the subset of these that occurred at a copy number of at least 10, and assembled them using SPAdes^73^ with “--only-assembler” and “--sc”. This process yielded the sequences of two candidate insertion sequence elements within the *B. caccae* genome.

### Insertion sequence abundance estimation

Illumina Truseq short-read data were aligned with BWA^61^ to the insertion sequence integration regions obtained from read cloud and Athena assembly of the clinical microbiome data. Reads recruited to each insertion locus were realigned with STAR^74^ in order to obtain gapped alignments spanning the insertion sequence. Gapped alignments, representing the genome sequence prior to insertion (“ancestral strain”), were counted for each insertion. Ancestral strain fraction was expressed as the number of observed gapped alignments divided by the median sequence coverage within the neighboring 10kb of sequence.

### Bacteroides caccae isolation

Members of the *Bacteroides fragilis* group, including *Bacteroides caccae*, were isolated directly from stool by streaking stool on solid BHI medium (containing 37g/L brain heart infusion powder, 1% v/v Remel defibrinated sheep blood, 100ug/mL gentamicin, and 1.5% agar) under anaerobic conditions in a Bactron 300 anaerobic chamber (Sheldon Manufacturing Inc, Cornelius, OR). Individual colonies matching the described *B. fragilis* group morphology of circular, entire, convex, gray, translucent, shiny and smooth^75^ were picked into 5mL liquid tryptone yeast glucose (TYG) media^76^ and incubated overnight inside the anaerobic chamber at 37°C. Aliquots of liquid cultures with glycerol added to 30% final concentration were frozen at −80°C. Subsequent whole genome sequencing of DNA extracted from these liquid cultures was used to confirm the species identity of each isolate.

### Isolate sequencing, assembly, and annotation

We extracted DNA using the Qiagen Gentra Puregene Bacteria DNA kit and prepared Illumina Nextera XT short-read sequencing libraries from all of the isolates that were cultured from the stool samples. The resulting 53 sequencing libraries were multiplexed and sequenced on a single Illumina HiSeq 4000 lane (Supplementary Table 8).

Short reads from each library were trimmed using the same procedure applied to stool sample libraries. Trimmed reads were then assembled using SPAdes^73^ to obtain genome drafts, and contigs from each draft were taxonomically annotated using Kraken v0.10.6^69^ with a custom database constructed from the Refseq and Genbank^70,71^. Genes were identified using Prokka v1.12 ^65^.

### Genotyping of IS614 and genomic island integration loci in *B. caccae* isolates

Alignments of the short reads back to their respective genome drafts allowed discovery of IS614 integration loci present in each *B. caccae* isolate genome. For each isolate, a modified genome draft was first obtained by masking sequences of partially reconstructed IS614 elements, and inserting a single fully assembled IS614 sequence. Raw reads were then mapped back to this modified genome draft using BWA^77^. Candidate flanking sequences downstream of the IS614 promoter in each isolate were then found by examination of spanning read pairs mapping between the added IS614 sequence and other sequence contigs. Isolates share flanking sequences as the majority of integrations occur more than once. These were reconciled to obtain a unique flanking set and the IS614 genotype was determined for all isolates from this unique set.

In order to search for potential large genomic island sequences that are exclusively integrated into *B. caccae* isolate genomes of a particular time point, *k*-mer counts from all isolate genome drafts are first obtained using Jellyfish^72^ with k = 31. For pairs of time points (A, C) and (C, D), we then searched for sets of *k*-mers that were exclusively present in the antecedent points, but not preceding ones. Each set of *k*-mers was then assembled them using SPAdes^73^ with “--only-assembler” and “--sc” to obtain candidate island sequences. This procedure yielded only two such large genomic islands, both of which were also found using comparisons of our read cloud *B. caccae* drafts.

### Antibiotic susceptibility testing and MIC determination

*B. caccae* isolate strains were first selected for testing on the basis of their IS614 genotype determined by whole genome sequencing. One isolate from time point A (A2) was determined to contain both *B. caccae* and another *Bacteroides* species most closely related to *Bacteroides uniformis*. A2 was subsequently re-streaked on BHI plates that allowed isolation of the *B. caccae* strain, which was verified using PCR (described below). A new glycerol stock containing only *B. caccae* was prepared for testing.

Ciprofloxacin and trimethoprim susceptibility was assessed using a broth microdilution MIC method adapted from CLSI^78^ as follows. Culture tubes, TYG media, 96-well assay plates, pipette tips and all other labware were reduced overnight in an anaerobic chamber prior to culture. Each strain was first grown directly from prepared glycerol stocks in 2.5mL of TYG liquid culture for 48 hours to log-phase. A 25mg/mL drug stock of ciprofloxacin was dissolved in acidified water (0.1M HCl) and a 50mg/mL drug stock of trimethoprim was dissolved in DMSO. Both antibiotics stock solutions were reduced overnight prior to testing. 96-well flat-bottom assay plates were setup to have final reaction volumes of 200μL with one control well and 11 wells with two-fold serial dilutions of each drug. Plates were prepared with TYG and drug stocks such that wells with the highest drug concentrations would contain a final concentration of 4096mg/μL and 512mg/μL for ciprofloxacin and trimethoprim, respectively. OD6OO readings of each liquid culture were performed prior to assay plate setup and used to resuspend inocula in fresh TYG media to normalize turbidity across the tested strains. Wells in the assay plates were then inoculated with 100μL of resuspended inocula such that the final turbidity was a 1:200 dilution of a 0.5 McFarland standard (OD600 of roughly 0.1). Each selected *B. caccae* isolate strain was tested using at least two and up to four replicates. Culture plates were incubated for 48 hours at 37°C in the anaerobic chamber. OD600 was then measured for each plate on a BioTek Epoch spectrophotometer (BioTek Instruments, Inc, Winooski, VT). Prior to plate reading, wells were mixed by pipetting up and down to allow a more even measurement as all *B. caccae* strains were observed to flocculate over the incubation period.

Ciprofloxacin precipitated out of solution at concentrations higher than 512ug/mL after the 48 incubation period and these readings were excluded from further susceptibility analysis.

The area-under-the-curve (AUC) of each resulting OD600 growth curve above a blank reading of empty TYG media was used to determine an MIC of each tested replicate. The MIC was determined to be the highest drug concentration to contain ≥ 90% of all growth as measured by AUC.

### PCR amplification

PCR was performed to verify insertion sequence presence across time points at select loci where they were found to be assembled. PCR reactions contained Phusion High-Fidelity DNA Polymerase (New England BioLabs, Ipswich, MA) with Phusion HF Buffer and NEB Deoxynucleotide Solution Mix. Primers were obtained from Elim Biopharm (Hayward, CA) with target melting temperature of 60°C. Reactions consisted of 6μL 5X Phusion HF buffer, 0.6μL 10mM dNTP, 0.3μL Phusion, 2μL 10mM forward primer, 2μL 10mM reverse primer, 1μL template DNA, and PCR clean water to 30μL. Thermocycling was performed with 30 seconds of denaturation at 98°C followed by 35 cycles of 5 seconds of denaturation at 98°C, 10 seconds of annealing at 65°C, and 30 seconds per kilobase extension at 72°C. This was followed by final extension at 72°C for 5 minutes and an indefinite hold at 4°C.

## Code availability

The Athena assembler together with a demonstration dataset can be found at https://αithub.com/abishara/athena_meta. The workflows used to produce and evaluate bins and generate the circos genome plot and composition barplot can be found at https://αithub.com/elimoss/metagenomics_workflows.

## Data availability

The datasets generated during the current study are available in the NCBI Sequence Read Archive under Bioproject accession PRJNA434731.

## Acknowledgements

The authors would like to thank Alexandra Sockell for assistance operating the NextSeq 500, Harmony Folse for participation in the isolation and culture of various *Bacteroides caccae* strains, and Arend Sidow for valuable feedback on the manuscript. This work was supported by NCI K08 CA184420, the Amy Strelzer Manasevit Award from the National Marrow Donor Program, and a Damon Runyon Clinical Investigator Award to A.S.B. E.L.M. was supported by National Science Foundation Graduate Research Fellowship DGE-114747. A.B. was supported by the Stanford Genome Training Program (SGTP; NIH/NHGRI), the Training Grant of the Joint Initiative for Metrology in Biology (JIMB; NIST), and the Stanford School of Medicine Dean’s Postdoctoral Fellowship. Access to shared compute resources was supported in part by NIH P30 CA124435 using the Stanford Cancer Institute Shared Resource Genetics Bioinformatics Service Center.

## Author contributions

E.L.M., A.B., A.S.B. and S.B. conceived of the study. E.L.M., C.H., C.W., Z.W. and H.J. prepared read cloud libraries. E.L.M., E.T., J.K., and T.A. collected samples, extracted DNA and RNA and prepared short-read sequencing libraries. E.L.M. designed and selected samples, and performed read cloud sequencing, PCR validation, and Sanger sequencing. A.B. and S.B. conceived of the assembly approach. A.B. implemented the Athena assembler. A.B., E.L.M, R.N.C., and S.Z. carried out the microbiological experiments. E.L.M. and A.B. carried out all analyses, wrote the manuscript, and generated figures. All authors commented on the manuscript.

